# AGL16 negatively modulates stress response to balance with growth

**DOI:** 10.1101/2021.02.16.431464

**Authors:** Ping-Xia Zhao, Jing Zhang, Si-Yan Chen, Jie Wu, Jing-Qiu Xia, Liang-Qi Sun, Shi-Song Ma, Cheng-Bin Xiang

## Abstract

Sensile plants constantly experience environmental stresses in nature. They must have evolved effective mechanisms to balance growth with stress response. Here we report the MADS-box transcription factor AGL16 acting as a negative regulator in stress response. Loss-of-*AGL16* confers resistance to salt stress in seed germination, root elongation, and soil-grown plants, while elevated *AGL16* expression confers the opposite phenotypes compared with wild type. However, the sensitivity to ABA in seed germination is inversely correlated with *AGL16* expression level. Transcriptomic comparison revealed that the improved salt resistance of *agl16* mutant was largely attributed to enhanced expression of stress responsive transcriptional factors and genes involved in ABA signaling and ion homeostasis. We further demonstrated that AGL16 directly binds to the CArG motifs in the promoter of *HKT1;1*, *HsfA6a,* and *MYB102* and represses their expression. Genetic analyses with double mutants also support that *HsfA6a* and *MYB102* are target genes of AGL16. Taken together, our results show that AGL16 acts as negative regulator transcriptionally suppressing key components in stress response and may play a critical role in balancing stress response with growth.

## INTRODUCTION

Plants suffer abiotic stresses in their life cycle. Soil salinity is a major abiotic stress that induces osmotic and oxidative stress and disturbs cellular ion and redox homeostasis, resulting in a substantial decrease of crop productivity worldwide (Miller et al., 2010; Schroeder et al., 2013; Golldack et al., 2014).

Plants have evolved a sophisticated network to cope with unfavorable environmental stresses. When exposed to salt stress, plants have effective responses to sense both hyperosmotic change and ionic fluctuation to deal with these stresses (Deinlein et al., 2014). But little is known about how salt stress is sensed by plants. Potential candidate of Na^+^ sensors for salinity stress sensing in plants were predicted, including SOS1 Na^+^/H^+^ antiporters, histidine kinase AtHK1, cation-H^+^ exchangers, NCX Na^+^/Ca^2+^ exchangers, NSCC/NADPH oxidase, mechanosensory channels and transporters, cyclic nucleotide receptors, purino-receptors, annexins, and H^+^-ATPase/GORK tandem (Urao et al., 1999; Zhu, 2003; Kurusu et al., 2013; Maathuis, 2014; Shabala et al., 2015; Wu, 2018; Duszyn et al., 2019). For hyperosmotic sense, it may be closely coupled with Ca^2+^ channels as demonstrated by rapidly increased Ca^2+^ levels in cytosolic with NaCl or mannitol treatment (Knight et al., 1997; Kurusu et al., 2013; Yuan et al., 2014; Manishankar et al., 2018). To date, *Arabidopsis* reduced hyperosmolality-induced [Ca^2+^]_i_ increase 1 (OSCA1) functions as an osmosensor in plants, which was involved in osmotic stress induced fast signaling response by formed a hyperosmolality-gated calcium-permeable channel (Yuan et al., 2014). Recently, plant cell-surface GIPC sphingolipids was reported to sense salt to trigger Ca^2+^ influx (Jiang et al., 2019). Salt stress-induced long-distance Ca^2+^ waves rapidly participate in stress response in plants, which is devoted to whole-plant stress tolerance by depending on the activity of the ion channel protein Two Pore Channel 1 (TPC1) (Choi et al., 2014).

Na^+^ and Cl^−^ accumulate to high levels under salt stress, leading to physiological disorders in plants. Therefore, sustaining low Na^+^ level and Na^+^ exclusion in shoots are pivotal for decreasing the damage of glycophytes in salt-affected soil (Deinlein et al., 2014; Munns and Gilliham, 2015). The Salt-Overly Sensitive (SOS) pathway including SOS1, SOS2, and SOS3 and transporting Na^+^ across the plasma membrane has been well characterized (Zhu, 2002, 2016; Yang and Guo, 2018). The key plasma membrane Na^+^/H^+^ antiporter SOS1 transports Na^+^ in the root back into the soil solution, thus reducing Na^+^ load from the xylem sap, which somehow triggers a cytoplasmic Ca^2+^ signal (Qiu et al., 2002; Pardo et al., 2006; Zhu, 2016).

Tonoplast-localized Na^+^/H^+^ exchanger, NHX1 compartmentalized Na^+^ into the vacuole to avoid the cytosolic accumulation of toxic Na^+^ (Blumwald et al., 2000). Overexpression of *NHX1* increases salt tolerance in Arabidopsis transgenic plants (Apse et al., 1999). Moreover, NHX1 and NHX2 are essential for active K^+^ uptake at the tonoplast, pH regulation in vacuole, and turgor generation for cell expansion (Leidi et al., 2010; Bassil et al., 2011; Barragan et al., 2012).

Efficient Na^+^ retrieval from the xylem stream to the surrounding tissues is important to reduce the accumulation of Na^+^ in shoots. High-affinity K^+^ transporter (HKT) family members are involved in Na^+^-selective transport and Na^+^ accumulation in shoots by mediating xylem Na^+^ unloading as shown in several plant species (Maser et al., 2002; Ren et al., 2005; Sunarpi et al., 2005; Horie et al., 2009). Arabidopsis *HKT1;1* functions in Na^+^ retrieval from the xylem and K^+^ homeostasis in shoots during salinity stress (Maser et al., 2002; Sunarpi et al., 2005; Davenport et al., 2007). The Arabidopsis *hkt1;1* mutant is hypersensitive to salt stress due to over accumulation of Na^+^ in shoots (Maser et al., 2002; Berthomieu et al., 2003), while enhanced expression of *HKT1;1* decreased Na^+^ level in shoots under salt stress (Moller et al., 2009). HKT1 also positively contributes to natural selection of saline adaptive *Arabidopsis thaliana* accessions (An et al., 2017).

Hormone signals play crucial roles in transcriptional regulation in response to salinity (Geng et al., 2013). Salt stress causes rapid and dynamic changes in the expression of stress responsive genes in plants, which overlaps with the response to abscisic acid (ABA) (Fujita et al., 2011; Geng et al., 2013). Primary root and lateral root growth show divergent temporal dynamics during salt stress, in which ABA and GA signaling crosstalks (Duan et al., 2013).

Transcription factors (TF) are key components modulating stress adaptive pathways in plant stress responses. Core members of TFs, including basic leucine zipper (bZIP), MYB, MYC, (ERF)/APETALA2 (AP2), basic helix-loop-helix (bHLH), NAC, and WRKY family, are differentially expressed in response to exogenous salinity (Golldack et al., 2011; Deinlein et al., 2014). TFs mediate abiotic stress tolerance mostly by regulating stress-specific genes involved in ion homeostasis or osmotic adjustment.

MADS-box family members are known for their roles in the regulation of flowering development (Messenguy and Dubois, 2003; Pinyopich et al., 2003; Lee and Lee, 2010; Callens et al., 2018). Some members of this TF family function in seed germination and root development. For example, ANR1 regulates lateral root growth in response to change in the external NO_3_^-^ supply, affects primary root development (Zhang and Forde, 1998; Gan et al., 2005; Gan et al., 2012), and acts as a modulator synergistically with AGL21 to regulate the expression of *ABI3* in seed germination (Lin et al., 2019). XAL1/AGL12 acts as an important regulator of cell proliferation in the root (Rounsley et al., 1995; Tapia-Lopez et al., 2008; Garcia-Cruz et al., 2016). Furthermore, XAL2/AGL14 modulates auxin transport during Arabidopsis primary root development by regulating PIN1 and PIN4 expression (Garay-Arroyo et al., 2013; Alvarez-Buylla et al., 2019). Moreover, AGL21, a member of the ANR1 clade, positively regulates auxin accumulation in lateral root primordia and lateral root initiation and growth (Yu et al., 2014b).

In this work, we continue to dissect the molecular mechanisms of *EDT1/HDG11*-conferred stress resistance by focusing on the down-regulated *AGL16* in *edt1* mutant. Our study shows that *AGL16* is responsive to salt, osmotic stress, and ABA at the transcription level. Loss-of-function mutants *agl16-2* and *agl16* are more resistant to salt stress in seed germination and primary root elongation than the wild type, and more sensitive to ABA, indicating that AGL16 plays an important role in regulating salt tolerance in Arabidopsis. Transcriptomic analyses show that AGL16 negatively regulates the expression of stress responsive genes. We further demonstrated that AGL16 represses the transcription of *HKT1;1*, *HsfA6a*, and *MYB102* by directly binding to their promoters, suggesting that *HKT1;1*, *HsfA6a* and *MYB102* are target genes of AGL16, which is consistent with the genetic analyses of double mutants. Together, our study revealed that AGL16 acts as an important negative regulator in stress response and may participate in the balance between growth and stress response.

## RESULTS

### AGL16 negatively regulates seed germination in response to abiotic stress

To investigate the role of *AGL16* in stress response, we obtained a T-DNA insertion line that we named *agl16*, the CRISPR/Cas9-edited *AGL16* knockout mutant *agl16-2*, and generated *AGL16* overexpression lines (OX) as described (Zhao et al., 2020), then seed germination assays were conducted. The seeds of wild type (Col-0), *agl16-2*, *agl16*, OX-17, OX-24 lines were germinated on MS medium with or without salt, mannitol, or ABA, and germination % was checked every day for 7 days. On MS medium, the seed germination rates showed no obvious difference among the genotypes. However, on medium with salt or mannitol, *agl16-2* and *agl16* mutants seed germination were less sensitive compared with the wild type, while OX lines were more sensitive to the treatments. In response to ABA treatment, *agl16-2* and *agl16* mutants displayed a reduced germination rate compared with the wild type, while OX lines showed more resistant germination to ABA inhibition than Col-0 (Figure 1A-B). Additionally, green cotyledon percentages of these genotypes displayed similar responses to the treatments as their seed germination did (Figure 1C). These results indicate that AGL16 negatively regulates seed germination in response to abiotic stress.

**Figure 1.**
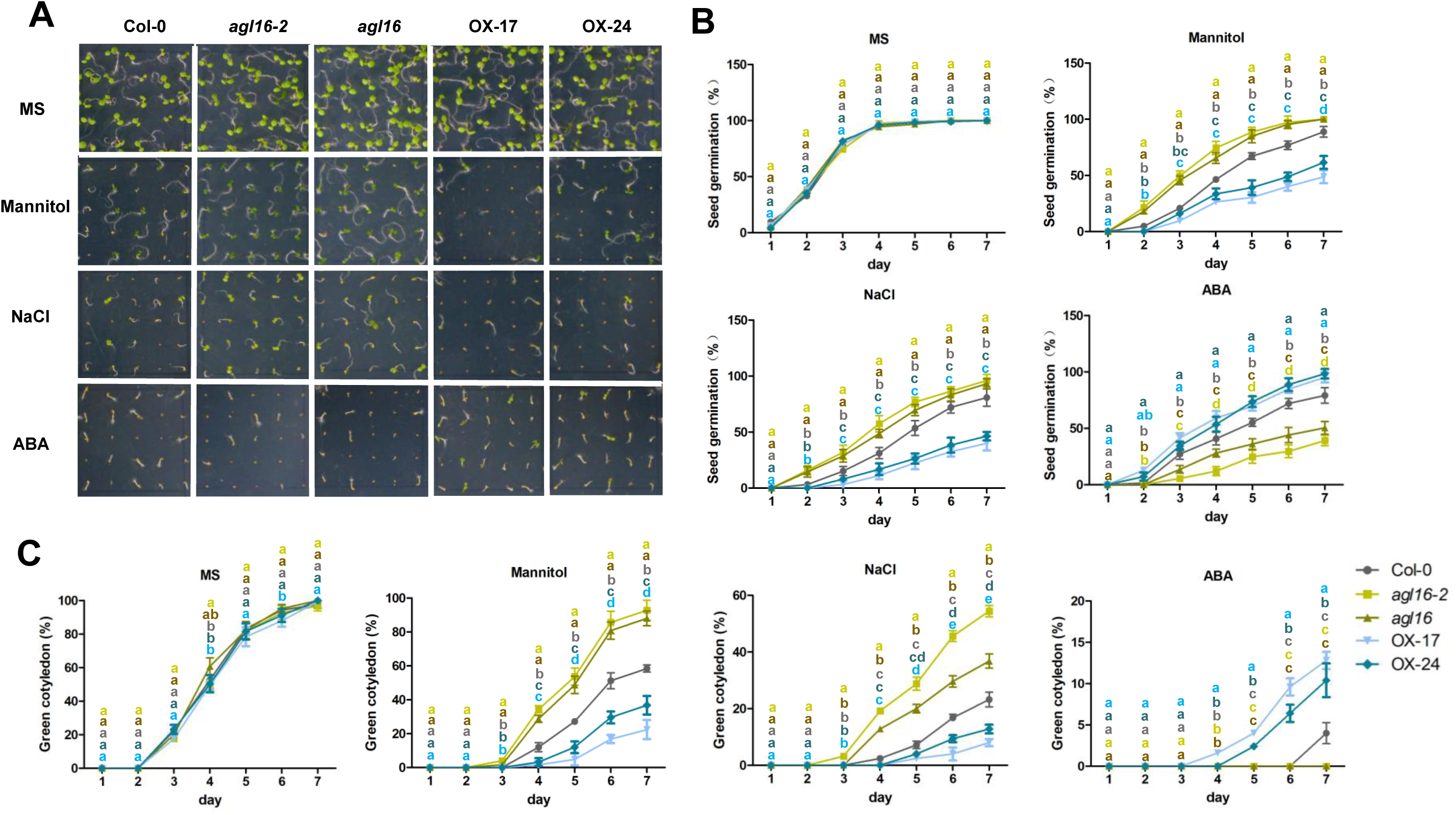
Response of *AGL16* to abiotic stress in seed germination. (A-C) Seed germination of Col-0, *agl16-2*, *agl16,* and *AGL16* overexpression (OX) lines. Seeds were horizontally germinated on MS medium with or without 120 mM NaCl, 250 mM mannitol and 0.5 µM ABA for 7 days. Photographs were taken (A). Seed germination rate was measured at the indicated time points (B). Green cotyledon rate was measured every day for 7 days (C). Values are mean ± SD (n=3 replicates, 60 seeds/replicate). Different letters indicate significant difference by one-way ANOVA (P < 0.05).

### AGL16 negatively regulates primary root elongation in response to abiotic stress

To confirm the role of AGL16 in stress response, we further conducted primary root elongation assays. The seeds of Col-0, *agl16-2*, *agl16*, OX-17, OX-24 lines were germinated on MS medium for 4 days, and then transferred to MS medium with or without salt, mannitol or ABA and vertically grew for another 8 days. Primary root length was measured at the indicated time points. Under normal growth conditions (MS), no difference was found in primary root length among Col-0, *agl16-2*, *agl16*, OX-17, OX-24 lines. When subjected to salt and mannitol treatment, *agl16-2* and *agl16* mutants showed a less sensitive primary root elongation, while the overexpression lines displayed a considerable reduction compared to Col-0 (Figure 2A-B). Similarly, the fresh weight of *agl16-2* and *agl16* mutants were significantly higher than that of Col-0 after salt and mannitol treatment, while OX lines exhibited lower fresh weight (Figure 2C). In response to ABA treatment, *agl16-2* and *agl16* mutant were more sensitive in primary root elongation, while the OX lines were much less sensitive compared to Col-0 (Figure 2A-B). Similarly, fresh weight was significantly decreased in *agl16-2* and *agl16* mutant compared with that of Col-0, while OX lines exhibited the opposite (Figure 2A-C).

**Figure 2.**
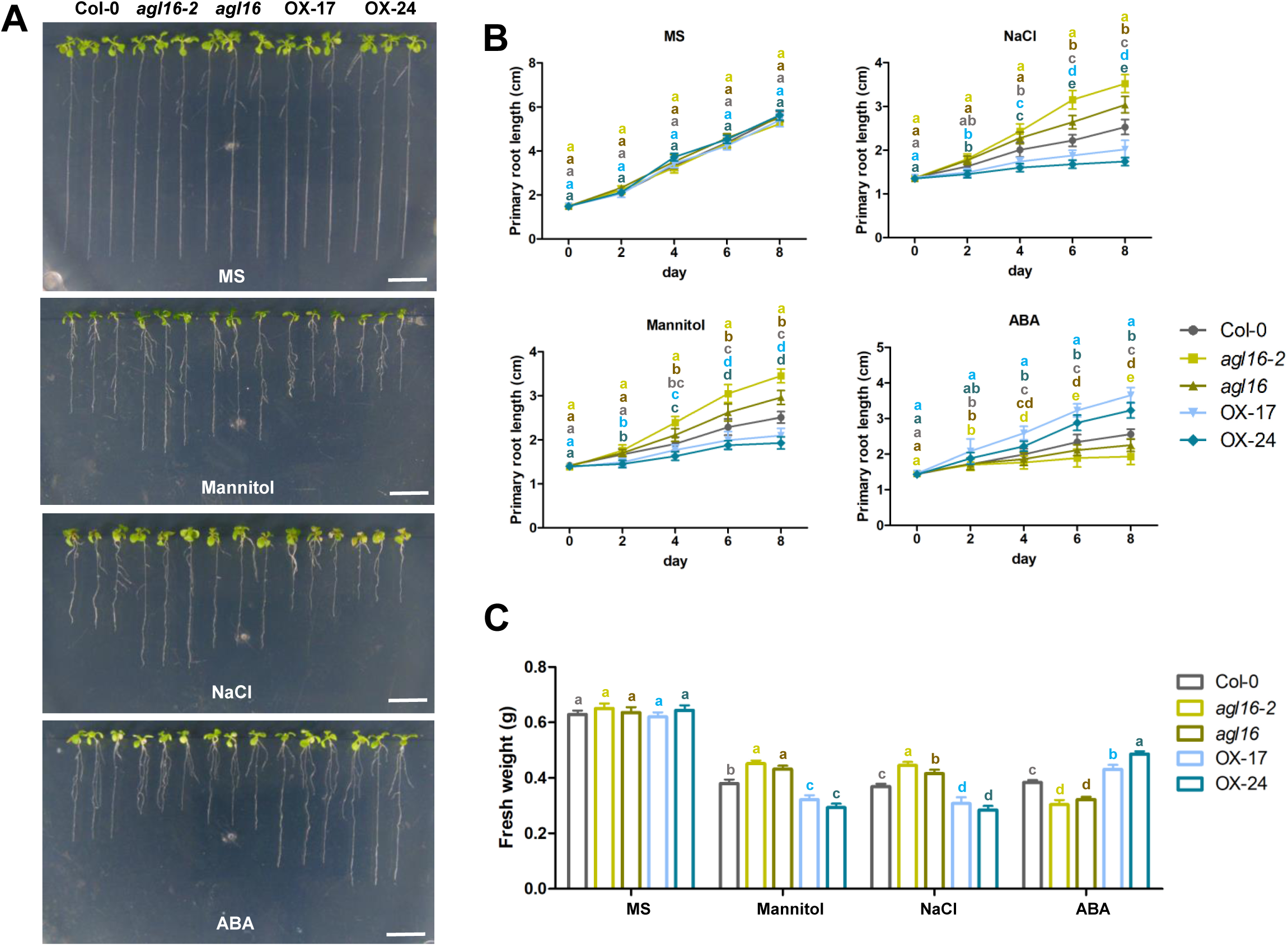
Primary root elongation of *agl16* mutants and *AGL16* overexpression lines in response to abiotic stress. (A-B) Primary root elongation. Seeds of Col-0, *agl16-2*, *agl16,* and OX lines were germinated on MS medium for 4 days respectively, then seedlings were transferred to MS medium with or without 120 mM NaCl, 250 mM mannitol and 5 µM ABA for 8 days. Photographs were taken (A) and primary root length was measured at the indicated time points (B). Values are the mean ± SD (n=3 replicates, 30 seedlings/replicate). Different letters indicate significant difference by one-way ANOVA (P < 0.05). (C) The fresh weights. The fresh weights of 12-day-old seedlings of Col-0, *agl16-2*, *agl16,* and OX lines were measured. Values are mean ± SD (n=3 replicates, 30 seedlings/replicate). Different letters indicate significant difference by one-way ANOVA (P < 0.05).

To explore whether cell division was affected by AGL16, we crossed the *agl16-2*, *agl16* mutant or OX lines with *CyclinB1;1-GUS* reporter, respectively to examine the mitotic activity in the root meristems. Under normal conditions, GUS signals were similar in the root meristem between the wild type, *agl16-2*, *agl16* mutant, and OX lines. In the salt and mannitol stressed roots, the activity of cell division was stronger in *agl16-2* and *agl16* mutants but significantly decreased in root meristem of OX lines compare with that of Col-0 (Figure 3). When treated with ABA, GUS signals were attenuated in *agl16-2* and *agl16* mutants compared to Col-0, while enhanced in the OX lines. Thus, mitotic activity in root meristem is one contributor to the AGL16-mediated primary root elongation in response to abiotic stress.

**Figure 3.**
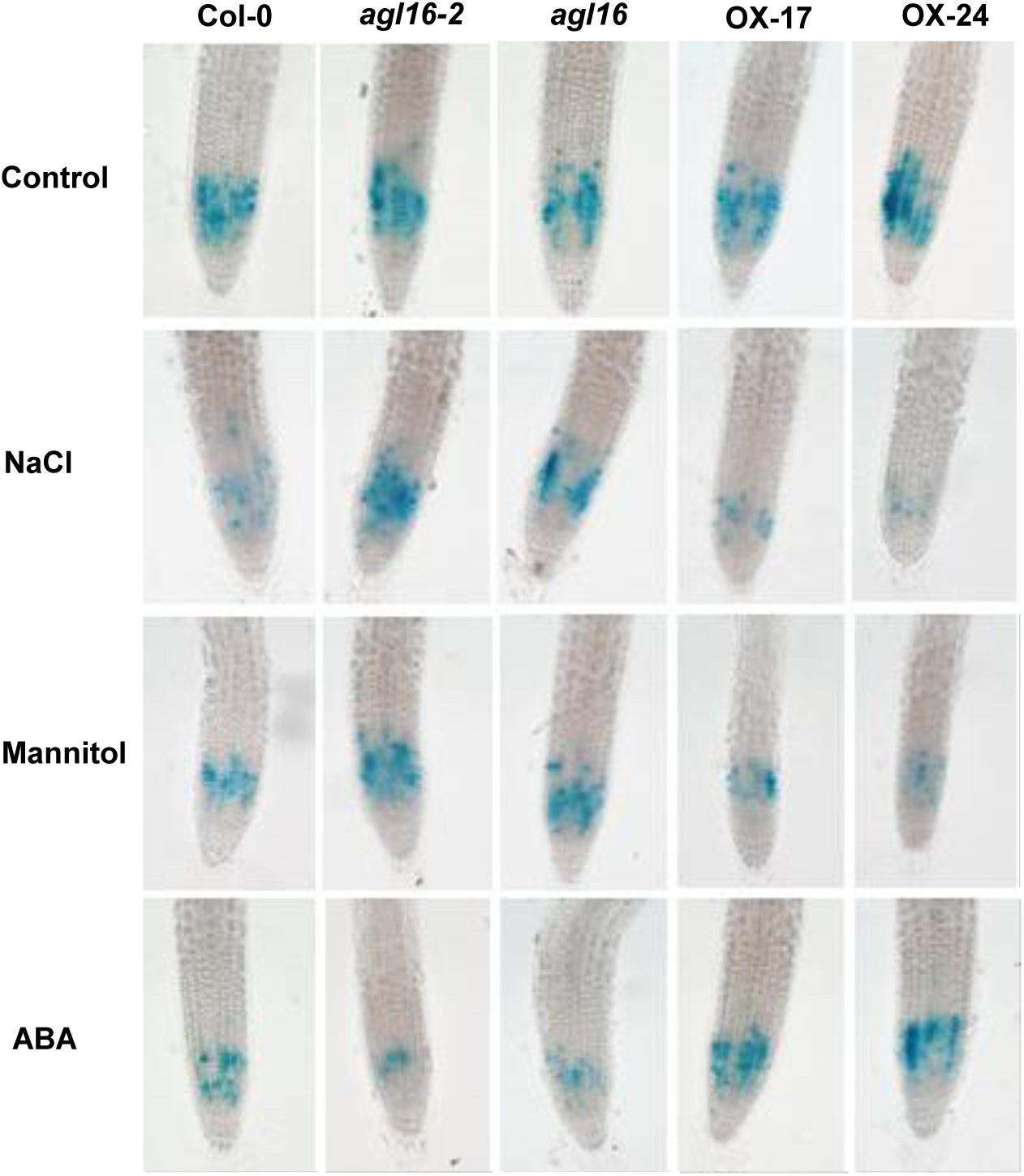
*CyclinB1;1* activity in primary root of Col-0, *agl16-2*, *agl16*, and *AGL16* overexpression lines. The *CyclinB1;1-GUS* maker was introduced in Col-0, *agl16-2*, *agl16*, and OX lines background by genetic crossing. The seeds were germinated on MS medium for 4 days, then seedlings were respectively transferred to MS medium with or without 120 mM NaCl, 250 mM mannitol, 5 µM ABA and grown for 3 days before GUS staining. Seedlings (>20) were incubated in GUS solution for 5 hours before photographs were taken. Bar=100 μm.

### AGL16 negatively regulates salt tolerance in Arabidopsis plants grown in soil

To evaluate the function of AGL16 in stress response of rosette stage plants, we performed salt tolerance assay on plants grown in soil. Soil-grown plants did not exhibit any apparent difference among Col-0, *agl16-2*, *agl16*, and OX lines under normal growth conditions. When 3-week old, soil-grown plants were watered with 250 mM NaCl solution and grown for 15 days before survival % was recorded. The loss-of-function mutants *agl16-2*, *agl16* lines exhibited a higher survival than Col-0, while OX-17 line showed the lowest survival (Figure 4A-B).

**Figure 4.**
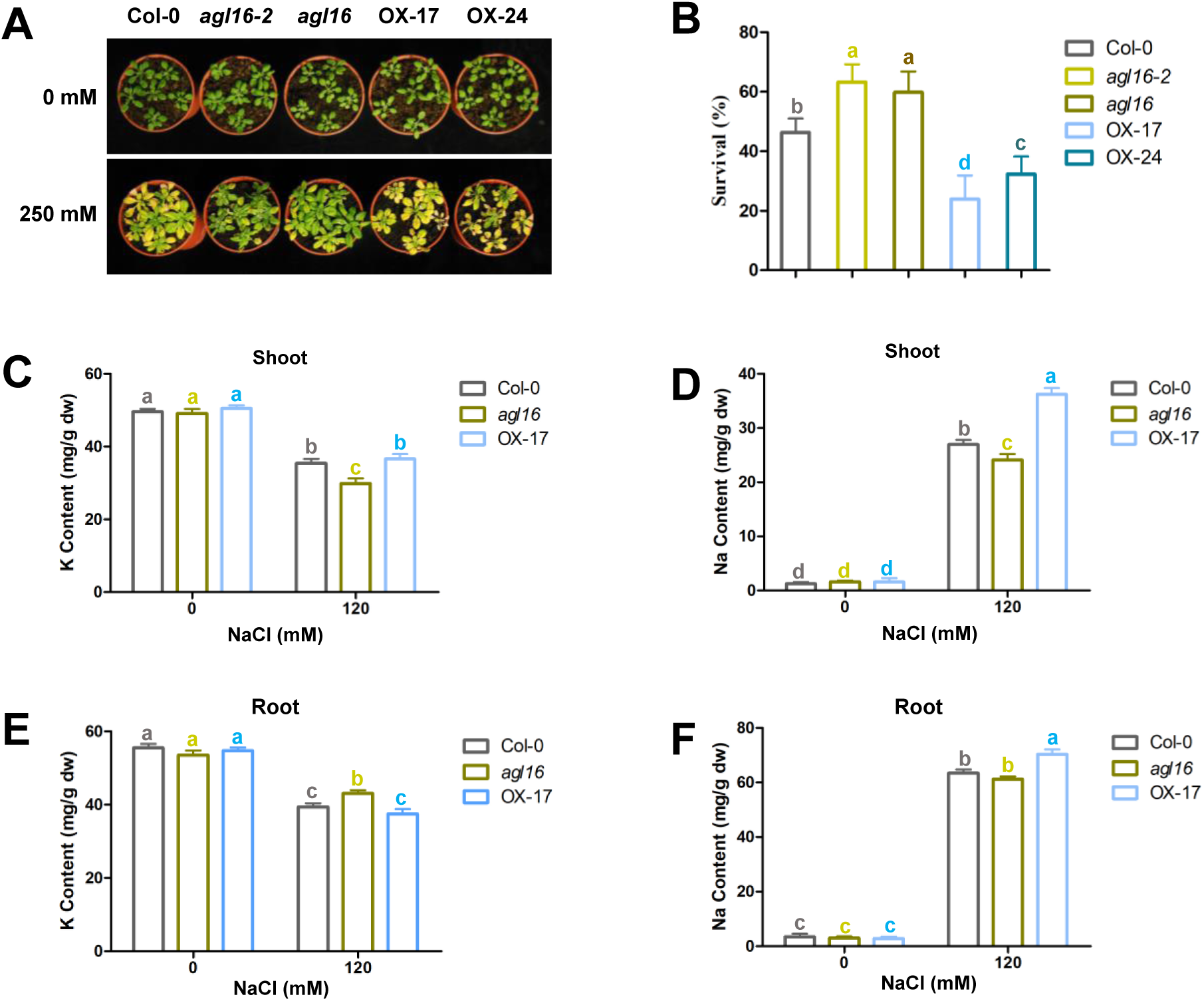
AGL16 negatively modulates salt tolerance in Arabidopsis plants grown in soil. (A-B) Salt tolerance assay in soil. 3-week old soil-grown seedlings of Col-0, *agl16-2*, *agl16*, and OX lines were irrigated with or without 250 mM NaCl for 2 weeks. Photographs were taken (A) and survival rate was calculated (B). Values are mean ± SD (n=3 replicates, 60 seedlings/replicate). Different letters indicate significant difference by one-way ANOVA (P < 0.05). (C-F) Ion accumulation in salt-treated Col-0, *agl16*, OX-17 plants. Seeds of Col-0, *agl16*, OX-17 lines were respectively germinated on MS medium for 4 days, then seedlings were transferred to MS medium with or without 120 mM NaCl and grown vertically for 10 days. Sodium, potassium content were quntified in shoots (C-D) and roots (E-F). Values are mean ± SD (n=3 replicates, 50 seedlings/replicate). Different letters indicate significant difference by one-way ANOVA (P < 0.05).

Next, we analyzed Na^+^ and K^+^ content in shoots and roots of Col-0, *agl16*, OX seedlings exposed to moderate salt stress. Under non-stress conditions, the content of Na^+^, K^+^ were similar in shoots and roots of Col-0, *agl16*, OX lines (Figure 4C-F). Under salt stress, less Na^+^ was accumulated in the shoots of *agl16* mutant than that of Col-0, while OX lines had significantly higher Na^+^ levels. However, in the roots, no significant difference in Na^+^ content was seen between Col-0 and *agl16* but the OX root had a higher level than Col-0 (Figure 4D, F). Under the same salt stress, shoot K^+^ levels did not differ between OX and Col-0 lines, but *agl16* mutant had lower K^+^ compared with that of Col-0 (Figure 4C). In contrast, root K^+^ level in *agl16* mutant was significantly higher than that of Col-0, while the OX had a similar K^+^ level as the wild type (Figure 4C, E). These results suggest that AGL16 may influence Na^+^ and K^+^ content and therefore ion homeostasis in roots and shoots, leading to altered salt resistance seen in *agl16* and OX lines.

### *AGL16* responds to multiple stresses and AGL16 protein always localizes in the nucleus

We previously showed higher expression levels of AGL16 in leaves, stem, and siliques by quantitative RT-PCR analyses (Zhao et al., 2020). To learn a more detailed tissue expression pattern, we generated *AGL16pro-GUS* transgenic plants and examined that the transcript abundance of *AGL16* in different tissues by GUS staining under normal conditions (Figure S1A). *AGL16* was primarily expressed in radicle of germinating seeds (Figure S1Aa-b). In 5-day-, 10-day-, 15-day- and 20-day-old plants, GUS staining was found in both leaves and roots (Figure S1Ac-f) where it was mainly located in root tip and stele (Figure S1Ag-h). In maturing plants, *AGL16* was mainly expressed in rosette leaves, cauline leaves, flowers, and siliques (Figure S1Ai-k).

To further study the response of *AGL16* expression to abiotic stress during root development, we examined the transcript level of *AGL16* by qRT-PCR in 7-day-old wild type (Col-0) seedlings treated with salt, mannitol, and ABA for 0, 0.5, 1, 2, 3, 6, 12, 24 hours. Under normal condition, the transcript levels of *AGL16* did not show significant difference. However, the expression of *AGL16* was down regulated after salt, mannitol treatment for 24 hours, and gradually upregulated by ABA treatment and peaked at 3 hours after the treatment (Figure S1B). We also observed the responses in *AGL16pro-GUS* transgenic plants to salt, mannitol, ABA treatment. GUS signals in root tip and stele gradually declined by salt and mannitol treatment. However, to ABA treatment, GUS signal peaked at 3 hours after the treatment and declined in root tip and stele (Figure S1C). The GUS staining results are largely consistent with the qRT-PCR results (Figure S1B).

Moreover, *AGL16* responds to multiple exogenous signals, such as 1-aminocyclopropane-1-carboxylic acid (ACC), indole-3-acetic acid (IAA) during root development (Figure S1D).

To localize AGL16 protein in the cell, we generated *35S:AGL16-GFP* transgenic plants and found that the subcellular localization of AGL16 protein was in the nucleus of 7-day-old seedling roots by confocal laser scanning microscope (Figure S1E).

### The response of *AGL16* to ABA is directly mediated by ABA signaling and biosynthesis pathway in modulating seed germination

Since loss-of-function mutants *agl16-2*, *agl16* were hypersensitive to exogenous ABA in seed germination, we next investigated the role of *AGL16* in imbibed seeds in response to ABA. qRT-PCR analyses showed that the transcript level of *AGL16* was strongly elevated by exogenous ABA in wild-type (Col-0) seeds after 2-3 days imbibition (Figure 5A). We further tested whether *AGL16* is involved in the regulation of ABA signaling by examining *AGL16* transcript level in ABA insensitive (*abi*) mutants in the absence or presence of exogenous ABA. As shown in Figure 5B, *AGL16* expression was significantly induced by ABA treatment only in Col-0, *abi4-1,* and *abi5-7* mutants. However, the induced expression of *AGL16* was abolished in *abi1-1*, *abi2-2*, *abi3-8* mutants. Meanwhile, the *agl16 abi5* double mutant exhibited alleviated ABA hypersensitivity (Figure 4C-E). These data suggest that *AGL16* acts downstream of ABI1, ABI2 and ABI3 in ABA signaling to regulate seed germination.

**Figure 5.**
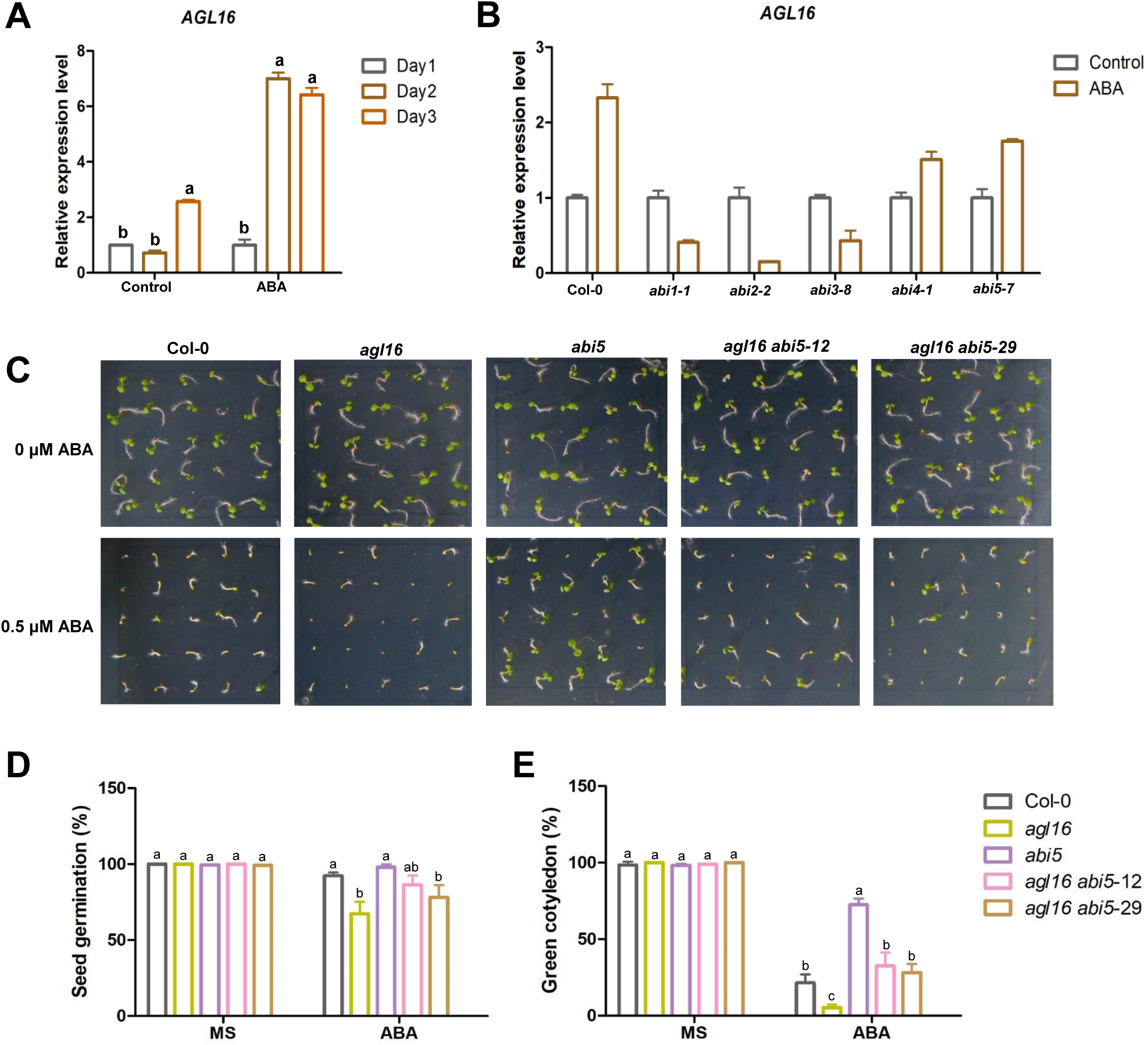
*AGL16* expression is dependent on ABA signaling in seed germination. (A) The response of *AGL16* expression to ABA in seeds. Vernalized wild type (Col-0) seeds were germinated on MS medium without (control) or with 0.5 µM ABA for 1, 2, 3 days. RNA was extracted from seeds and the transcript level of *AGL16* was analyzed by qRT-PCR. Values are mean ± SD (n=3 replicates). Different letters indicate significant difference by one-way ANOVA (P < 0.05). (B) Transcript levels of *AGL16* in *abi* mutants. Vernalized seeds of Col-0, *abi1-1*, *abi2-2*, *abi3-8*, *abi4-1*, and *abi5-7* were germinated on MS medium without (control) or with 0.5 µM ABA for 24 hours, respectively. RNA was extracted and the transcript level of *AGL16* was detected by qRT-PCR. Values are mean ± SD (n=3 replicates). (C-E) *AGL16* acts upstream of *ABI5*. Seeds of Col-0, *agl16*, *abi5*, *agl16 abi5* lines were horizontally germinated on MS medium with or without 0.5 µM ABA for 7 days. Photographs were taken (C). Seed germination rate (D) and green cotyledon rate were measured at 7 days after the end of stratification (E). Values are mean ± SD (n= 3 replicates, 60 seeds/ replicates). Different letters indicate significant difference by one-way ANOVA (P < 0.05).

**Figure 6.**
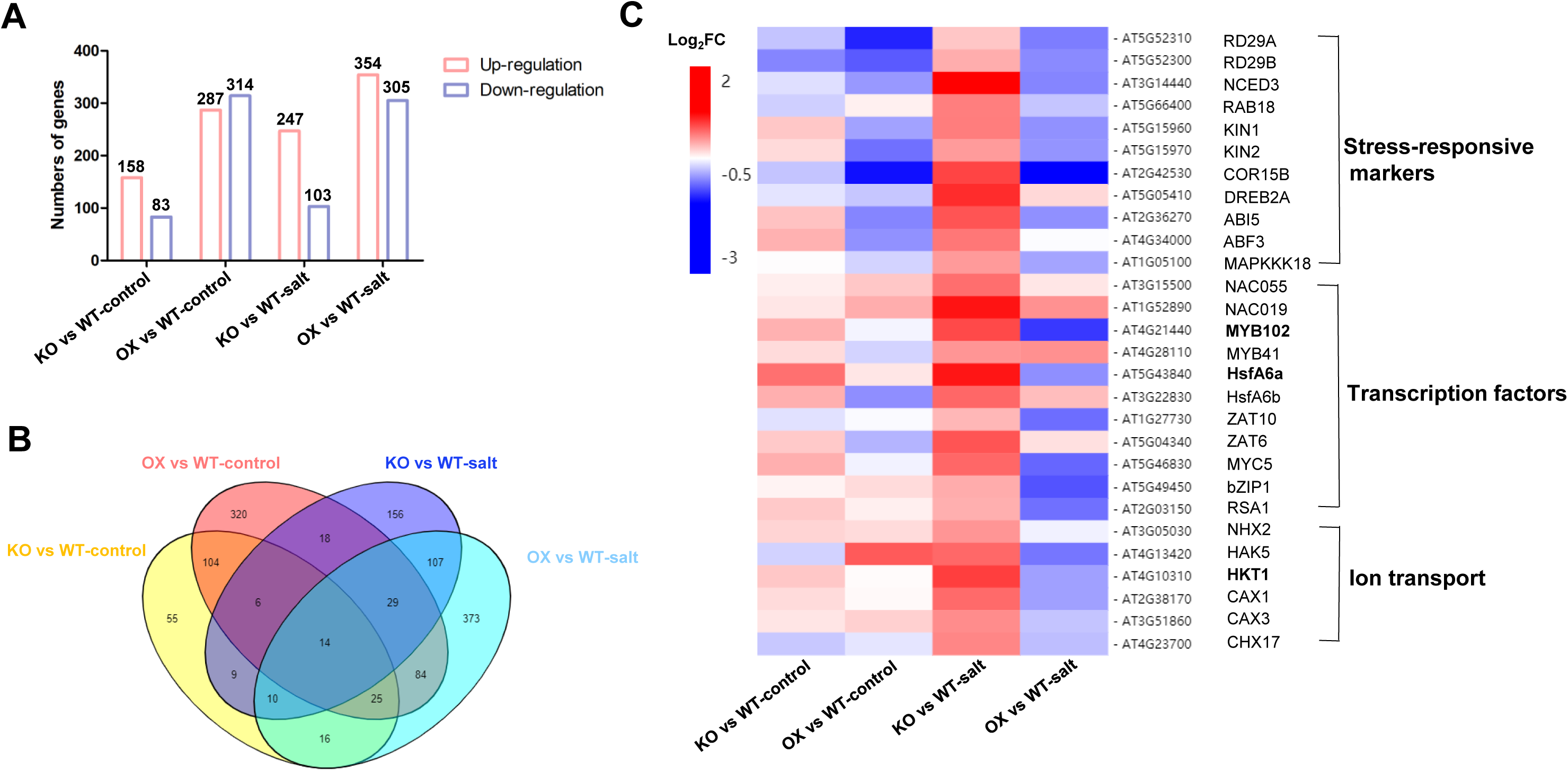
Transcriptomic analyses of differentially expressed genes (DEGs) affected by AGL16. (A) The number of differentially expressed genes (DEGs). The statistics data of differentially expressed genes in (KO vs WT)-control, (OX vs WT)-control and (KO vs WT)-salt, (OX vs WT)-salt groups. (B) Comparison of differentially expressed genes (DEGs) among (KO vs WT)-control, (OX vs WT)-control (KO vs WT)-salt, and (OX vs WT)-salt groups using Venn diagram. The numbers represent the total numbers of differentially expressed genes in different comparison groups. (C) Hierarchical clustering analysis of salt stress-related genes affected by AGL16 in DEGs. The heatmap represents fold changes in the abundance of gene transcripts in different comparison groups.

Furthermore, We examined the expression levels of genes associated with ABA signaling, including *ABI1*, *ABI2*, *SnRK2.2*, *SnRK2.3*, *ABI3*, *ABI4*, *ABI5*, and related to downstream ABA responsive genes, such as *EM1*, *EM6*, *RAB18*, *RD29B*. qRT-PCR results showed that *ABI4*, *ABI5*, *EM6*, *RAB18*, *RD29B* were upregulated in *agl16* mutant and downregulated in OX line compare to Col-0 in repsonse to exogenously applied ABA (Figure S2F-H, J-K), suggesting that AGL16 is a key node in ABA-response signaling pathway during seed germination. In addition, the expression level of ABA biosynthesis-related genes *ABA2*, *NCED3*, *AAO3* was also upregulated in *agl16* mutant compared with that of Col-0 after ABA treatment (Figure S2M, O-P). Whereas, the expression of key ABA catabolic enzyme family genes *CYP707A1*-*CYP707A4* have no significant difference (Figure S2Q-T). Taken together, these results suggest that AGL16-modulated seed germination involves ABA signaling and biosynthesis.

### Enhanced transcript levels of salt stress-responsive genes in *agl16* mutant

To understand how AGL16 regulates salt tolerance in Arabidopsis, we measured the expression levels of salt stress-related genes in the wild type, *agl16,* and OX seedlings in response to salt stress with qRT-PCR. The transcript levels of *RD29A*, *RD29B*, *RAB18*, *RD22,* and *CBF4* were significantly upregulated in *agl16* mutant compared with those in Col-0 plants (Figure S3A-E). The expression levels of *SOS1*, *SOS2*, *NHX1*, which are salt-inducible genes involved in ion homeostasis, were also altered in *agl16* mutant but not significantly changed compared with wild type (Figure S3F-H). In contrast, transcript level of *HKT1;1* was markedly elevated in *agl16* mutant but reduced in OX line (Figure S3I), suggesting that AGL16 may directly repress the expression of *HKT1;1* considering the *AGL16* expression pattern in root stele and leaf vasculature (Figure S1Ah) which is similar to Na^+^ transporter *HKT1;1* (Maser et al., 2002).

### RNA-seq analyses reveal global AGL16-regulated genes

To explore the regulatory network of AGL16-mediated stress response in plant growth, we performed RNA-seq analysis to identify differentially expressed genes (DEGs) (> 1.5 fold change, p < 0.05) between 7-day-old Col-0, *agl16*, and OX-17 seedlings exposed to salt treatment for 0, 3 hours. Figure S4A-D shows that the most of genes were concentrated, only a few of genes were scattered, indicating expression of these genes was strongly regulated by AGL16. Comparative analysis of the DEGs between *agl16* mutant, OX and Col-0 revealed that the number of DEGs was increased in (KO vs WT)-salt and (OX vs WT)-salt groups, suggesting that AGL16 influences the transcript profile in response to salt stress (Figure 5A). Furthermore, a small fraction of genes were differentially expressed between *agl16*, OX line, and Col-0 by comparison of DEGs number among (KO vs WT)-control, (OX vs WT)-control and (KO vs WT)-salt, (OX vs WT)-salt groups (Figure 5B). These co-regulated genes in *agl16*, OX and Col-0 line may be the candidate targets of AGL16.

In order to gain further insight into the regulatory network of AGL16 in response to short-term salt stress, a gene ontology (GO) analysis was performed for significantly up-regulated and down-regulated genes in (KO vs WT)-control, (OX vs WT)-control, (KO vs WT)-salt, and (OX vs WT)-salt pair-wise groups based on the biological process. Many DEGs from the (KO vs WT)-control group involved in pathways of response to heat, wounding, salt stress, jasmonic acid were highly enriched (Figure S5A). Several categories, including microtubule movement, response to wounding, jasmonic acid, heat, have preferential enrichment in (OX vs WT)-control group (Figure S5B). When treated with salt, numerous genes from the (KO vs WT)-salt group were assigned to abiotic stress-related terms, including response to water deprivation, salt stress, abscisic acid, and osmotic stress (Figure S5C). DEGs were significantly enriched for terms including glucosinolate biosynthesis, response to water deprivation, oxidation-reduction process, and DNA replication in (OX vs WT)-salt group (Figure S6D). GO analyses showed that salt-regulated genes were involved in AGL16-mediated abiotic tolerance.

Furthermore, the hierarchical clustering was conducted to analysis the expression profile of DEGs in response to salt stress. As shown in Figure 5C, the transcript levels of well-known abiotic stress- or ABA-responsive genes, such as *NCED3*, *RAB18*, *KIN1*, *COR15B*, *DREB2A*, *ABI5*, *ABF3* were significantly higher in the KO vs WT (salt stressed) group. Moreover, the expression level of salt stress-responsive transcription factors, including *NAC055*, *NAC019*, *MYB102*, *HsfA6a*, *ZAT6* were also higher in *agl16* mutant. Besides, AGL16 also affected the expression of a number of genes related to ion transport (Table S1).

Thus, we selected a few DEGs known to mediate abiotic stress response to verify their transcript levels in Col-0, *agl16*, OX-17 line. The transcript level of *HsfA6a* and *MYB102* were upregulated in *agl16* mutant and downregulated in OX line compared with the Col-0 under salt treatment for 3, 12 hours (Figure S6). However, the expression level of *bZIP1*, *NHX2* were not downregulated in OX compared with the Col-0 at 3, 12 hours. Therefore, we hypothesized AGL16 may directly regulate the expression of *HsfA6a* and *MYB102*, two known TFs in stress resistance (Denekamp and Smeekens, 2003; Hwang et al., 2014).

### AGL16 directly regulates *HKT1;1*, *HsfA6a* and *MYB102* in salt stress response

To determine whether AGL16 interacts directly with the promoter of *HKT1;1*, *HsfA6a* and *MYB102* or represses their expression indirectly, we found three putative AGL16-binding sites (CC-A/T-rich-GG or C-A/T-rich-G) in each 2.0 kb promoter region of *HKT1;1*, *HsfA6a*, *MYB102* (Figure 7-9A). We constructed transgenic Arabidopsis plants that overexpressed HA-AGL16 fusion protein, and performed chromatin immunoprecipitation (ChIP) assays. The promoter fragments (about 200 bp) with cis1 site of *HKT1;1*, cis1 and cis3 sites of *HsfA6a*, and cis2 site of *MYB102* were markedly enriched in *35S:HA-AGL16* transgenic plants by PCR and qRT-PCR (Figure 7-9B-C). To test the ability of AGL16 as a transcription repressor, we constructed the effector that the coding sequence of *AGL16* was driven by the cauliflower mosaic virus (CaMV) 35S promoter, and reporters that a 2000 bp promoter or 26 bp promoter fragments contained CArG motifs of *HKT1;1*, *HsfA6a* and *MYB102* were respectively fused to the luciferase (LUC) reporter gene. When effector and reporter were both transfected into Arabidopsis protoplasts, AGL16 was able to suppress the expression of the *HKT1;1*, *HsfA6a* and *MYB102* promoter-driven luciferase (LUC) reporter gene (Figure 7-9D), demonstrating that AGL16 can directly regulate the expression of *HKT1;1*, *HsfA6a* and *MYB102 in vivo*. Moreover, we performed yeast-one-hybrid (Y1H) and EMSA assays to confirm the specific binding. Y1H results displayed that AGL16 can interact with cis1 site of *HKT1;1* promoter, cis1 and cis3 sites of *HsfA6a* promoter, and cis2 site of *MYB102* promoter (Figure 7-9C). EMAS assays also demonstrated that AGL16 binds specifically to the cis element (Figure 7-9E). The above results suggest that AGL16 represses the expression of *HKT1;1*, *HsfA6a* and *MYB102* by directly binding to the CArG motifs in their promoters *in vivo* and *in vitro*.

**Figure 7.**
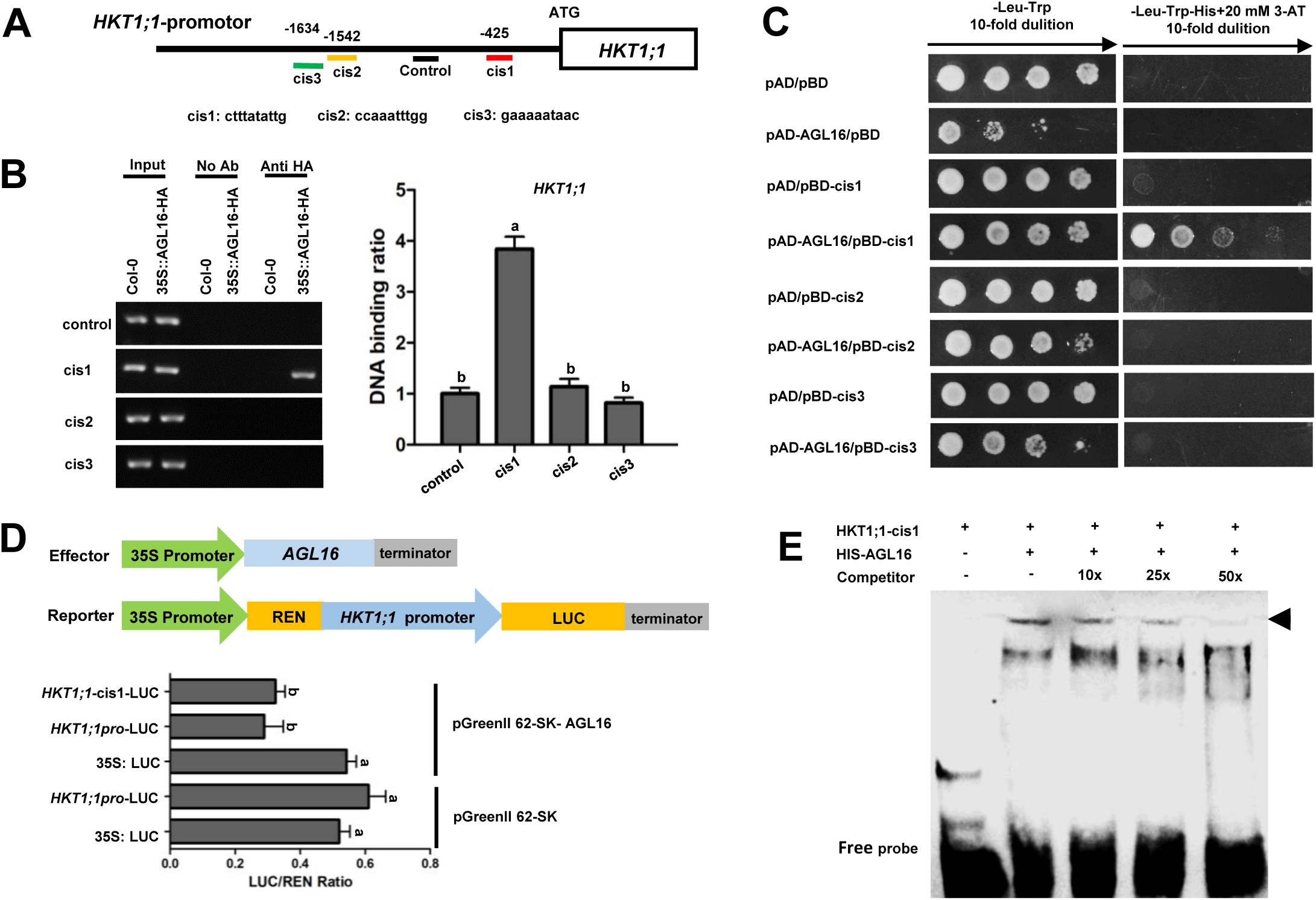
AGL16 binds to the CArG motif in the promoter of *HKT1;1*. (A) Schematic illustration of *HKT1;1* promoter with AGL16 candidate CArG motifs. The number represents the location of CArG motifs in *HKT1;1* promoter; Short lines of different colors mark the regions amplified in PCR and qRT-PCR analyses, while the black line represents the control region without CArG motifs. (B) ChIP-PCR assay. 7-day-old *35S:HA-AGL16* transgenic plants and wild type (Col-0) were used for the ChIP-PCR assay. A region of *HKT1;1* that does not contain CArG motifs was used as a control. About 200 bp fragment cis1 from *HKT1;1* promoter containing the CArG motif was enriched by anti-HA antibodies as shown in PCR and qRT-PCR analyses. Values are mean ± SD (n=3 replicates). Different letters indicate significant difference by one-way ANOVA (P < 0.05). (C) Yeast one-hybrid assay. The coding sequence of *AGL16* was constructed into pGADT7 (pAD) and the 26 bp fragment containing the CArG motif of *HKT1;1* promoter was cloned into pHIS2 (pBD). Different yeast culture dilutions (1:1, 1:10, 1:100, 1:1,000) were grown on SD/-Trp-Leu medium with or without His plus 20 mM 3-AT. pAD/pBD, pAD-AGL16/pBD, pAD/pBD- cis1, pAD/pBD-cis2, pAD/pBD-cis3 were used as negative controls. (D) Transient transactivation assays. Schematic illustration of the effector and reporters used in the transient transactivation assays. AGL16 effector were under the control of the CaMV 35S promoter. *HKT1;1* promoter and the 26 bp binding cis-element of *HKT1;1* promoter were respectively fused to the LUC gene as reporters. The firefly LUC and REN activities luciferase were detected by transient dual-luciferase reporter assays, and LUC/REN ratio was calculated. The REN activity was used as an internal control. Values are mean ± SD (n=3 replicates). Different letters indicate significant difference by one-way ANOVA (P < 0.05). (E) EMSA assay. The 26 bp cis-element containing the CArG motif of *HKT1;1* promoter was synthesized with biotin labelled at the 5’ end. Non-labelled probe was competitor. As indicated, AGL16-dependent mobility shifts were competed by the competitor probe in a dose-dependent manner.

**Figure 8.**
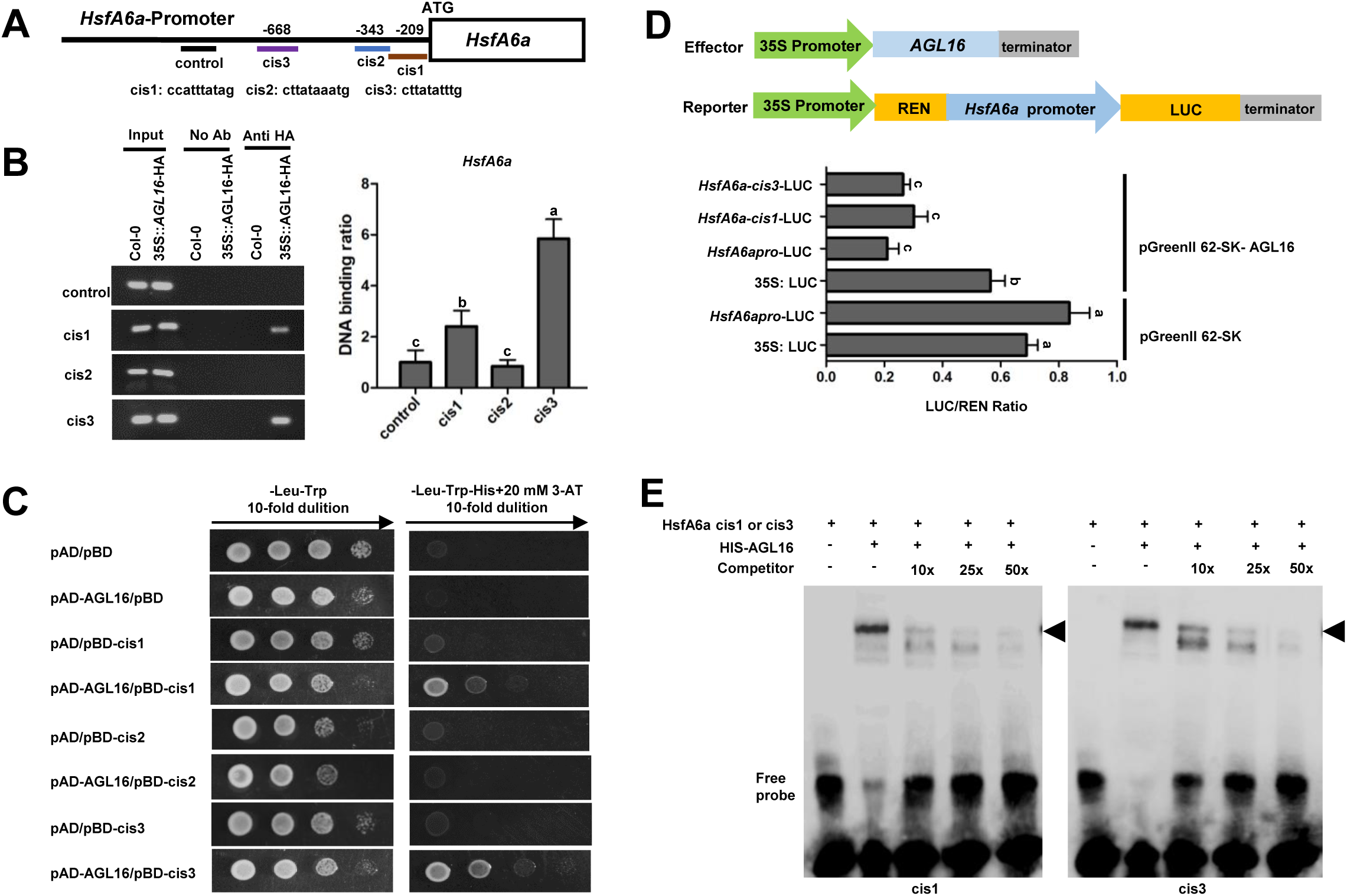
AGL16 binds to the CArG motifs in the promoter of *HsfA6a*. (A) Schematic illustration of *HsfA6a* promoter with candidate CArG motifs. The number represents the location of CArG motifs in *HsfA6a* promoter; Short lines of different colors mark the regions amplified in PCR and qRT-PCR analyses, while the black line represents the control region without CArG motifs. (B) ChIP-PCR assay. 7-day-old *35S:HA-AGL16* transgenic plants and wild type (Col-0) were used for the ChIP-PCR assay. A region of *HsfA6a* that does not contain CArG motifs was used as a control. About 200 bp fragment cis1 and cis3 from *HsfA6a* promoter containing CArG motifs were enriched by anti-HA antibodies as shown in PCR and qRT-PCR analyses. Values are mean ± SD (n=3 replicates). Different letters indicate significant difference by one-way ANOVA (P < 0.05). (C) Yeast one-hybrid assay. The coding sequence of *AGL16* was constructed into pGADT7 (pAD) and the 26 bp fragment containing the CArG motif of *HsfA6a* promoter was cloned into pHIS2 (pBD). Different yeast culture dilutions (1:1, 1:10, 1:100, 1:1,000) were grown on SD/-Trp-Leu medium with or without His plus 20 mM 3-AT. pAD/pBD, pAD-AGL16/pBD, pAD/pBD- cis1, pAD/pBD-cis2, pAD/pBD-cis3 were used as negative controls. (D) Transient transactivation assays. Schematic illustration of the effector and reporters used in the transient transactivation assays. AGL16 effector were under the control of the CaMV 35S promoter. *HsfA6a* promoter and the 26 bp cis-elements of *HsfA6a* promoter were respectively fused to the LUC gene as reporters. The firefly LUC and REN activities luciferase were detected by transient dual-luciferase reporter assays, and LUC/REN ratio was calculated. The REN activity was used as an internal control. Values are mean ± SD (n=3 replicate). Different letters indicate significant difference by one-way ANOVA (P < 0.05). (E) EMSA assay. The 26 bp cis-element containing CArG motif of *HsfA6a* promoter were synthesized with biotin labelled at the 5’ end. Non-labelled probe was used as competitor. As indicated, AGL16-dependent mobility shifts were competed by the competitor probe in a dose-dependent manner.

**Figure 9.**
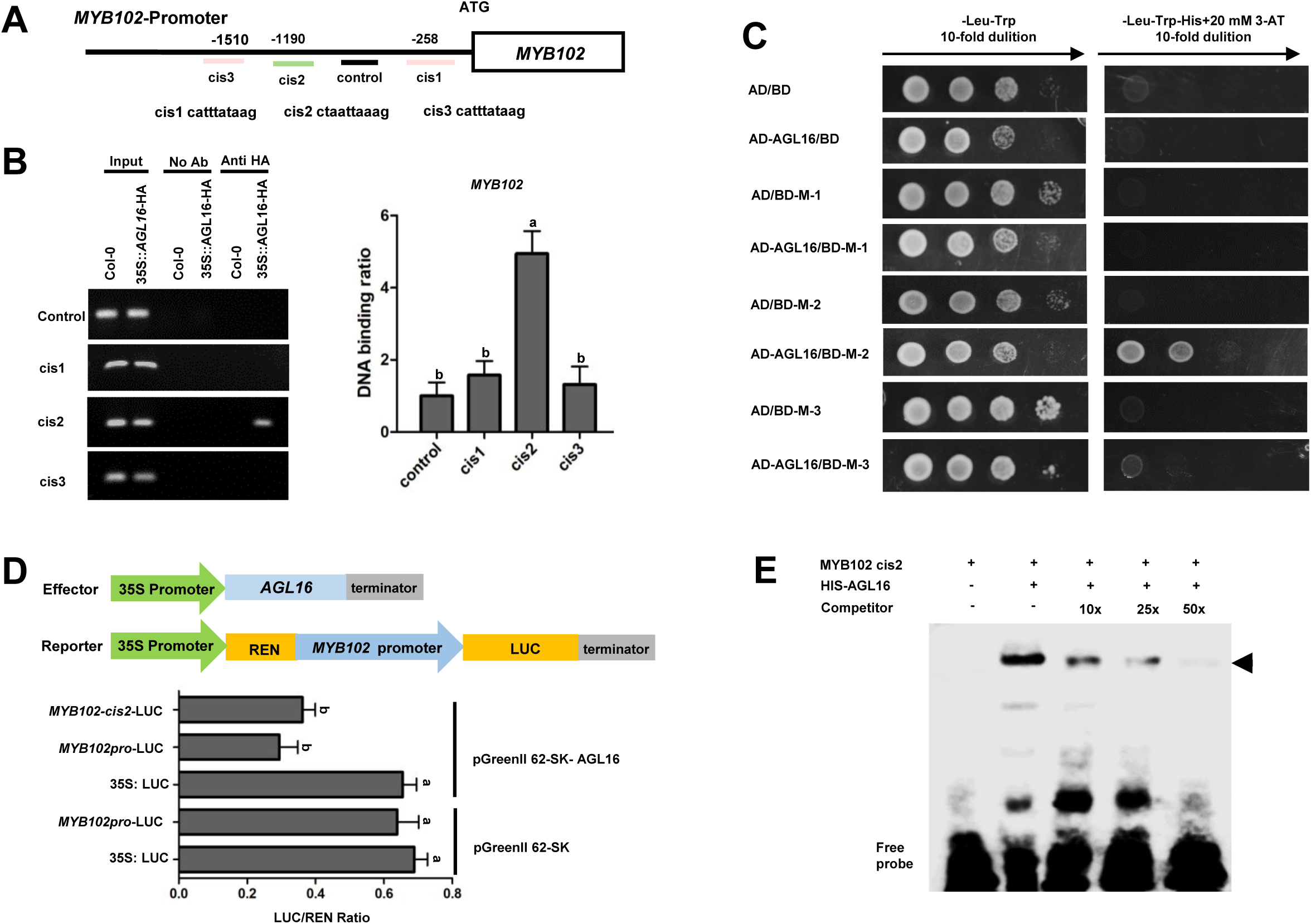
AGL16 binds to the CArG motifs in the promoter of *MYB102*. (A) Schematic illustration of *MYB102* promoter with AGL16 candidate CArG motifs. The number represents the location of CArG motifs in *MYB102* promoter; Short lines of different colors mark the regions amplified in PCR and qRT-PCR analyses, while the black line represents the control region without CArG motifs. (B) ChIP-PCR assay. 7-day-old *35S:HA-AGL16* transgenic plants and wild type (Col-0) were used for the ChIP-PCR assay. A region of *MYB102* that does not contain CArG motifs was used as a control. About 200 bp fragment cis2 from *MYB102* promoter containing CArG motifs was enriched by anti-HA antibodies as shown in PCR and qRT-PCR analyses. Values are mean ± SD (n=3 replicates). Different letters indicate significant difference by one-way ANOVA (P < 0.05). (C) Yeast one-hybrid assay. The coding sequence of *AGL16* was constructed into pGADT7 (pAD) and the 26 bp fragment containing CArG motif of *MYB102* promoter was cloned into pHIS2 (pBD). Different yeast culture dilutions (1:1, 1:10, 1:100, 1:1,000) were grown on SD/-Trp-Leu medium with or without His plus 20 mM 3-AT. pAD/pBD, pAD-AGL16/pBD, pAD/pBD-cis1, pAD/pBD- cis2, pAD/pBD-cis3 were used as negative controls. (D) Transient transactivation assays. Schematic illustration of the effector and reporters used in the transient transactivation assays. AGL16 effector were under the control of the CaMV 35S promoter. *MYB102* promoter and the 26 bp cis-element of *MYB102* promoter were respectively fused to the LUC gene as reporters. The firefly LUC and REN activities luciferase were detected by transient dual-luciferase reporter assays, and LUC/REN ratio was calculated. The REN activity was used as an internal control. Values are mean ± SD (n=3 replicates). Different letters indicate significant difference by one-way ANOVA (P < 0.05). (E) EMSA assay. The 26 bp cis-element containing CArG motifsof *MYB102* promoter was synthesized with biotin labelled at the 5’ end. Non-labelled probes was used as competitor. As indicated, AGL16-dependent mobility shifts were competed by the competitor probe in a dose-dependent manner.

### *HsfA6a* and *MYB102* acts downstream of *AGL16*

3*5S:HsfA6a* overexpression lines are insensitive to salt and drought, but sensitive to ABA (Hwang et al., 2014). To further verify the genetic interaction between *AGL16* and *HsfA6a, MYB102* in modulating seeds germination and primary root elongation in response to abiotic stress, we crossed *agl16* mutant with the *hsfa6a*, *myb102* mutants respectively and obtained the double mutants *agl16 hsfa6a* and *agl16 myb102*. The genetic analyses showed that seed germination rate and green cotyledon rate were similar in Col-0, *agl16*, *hsfa6a*, *agl16 hsfa6a* lines on MS medium (Figure 10A-C), but *agl16 hsfa6a* double mutant was more resistant to mannitol and NaCl in seed germination and post germination, but more sensitive to ABA than *hsfa6a* mutant (Figure 10A-C). As expected, the primary roots length of the *agl16 hsfa6a* double mutant was longer than those of *hsfa6a* mutant when subjected to salt and mannitol stress, while more inhibited than *hsfa6a* mutant by ABA treatment (Figure 10D-E).

**Figure 10.**
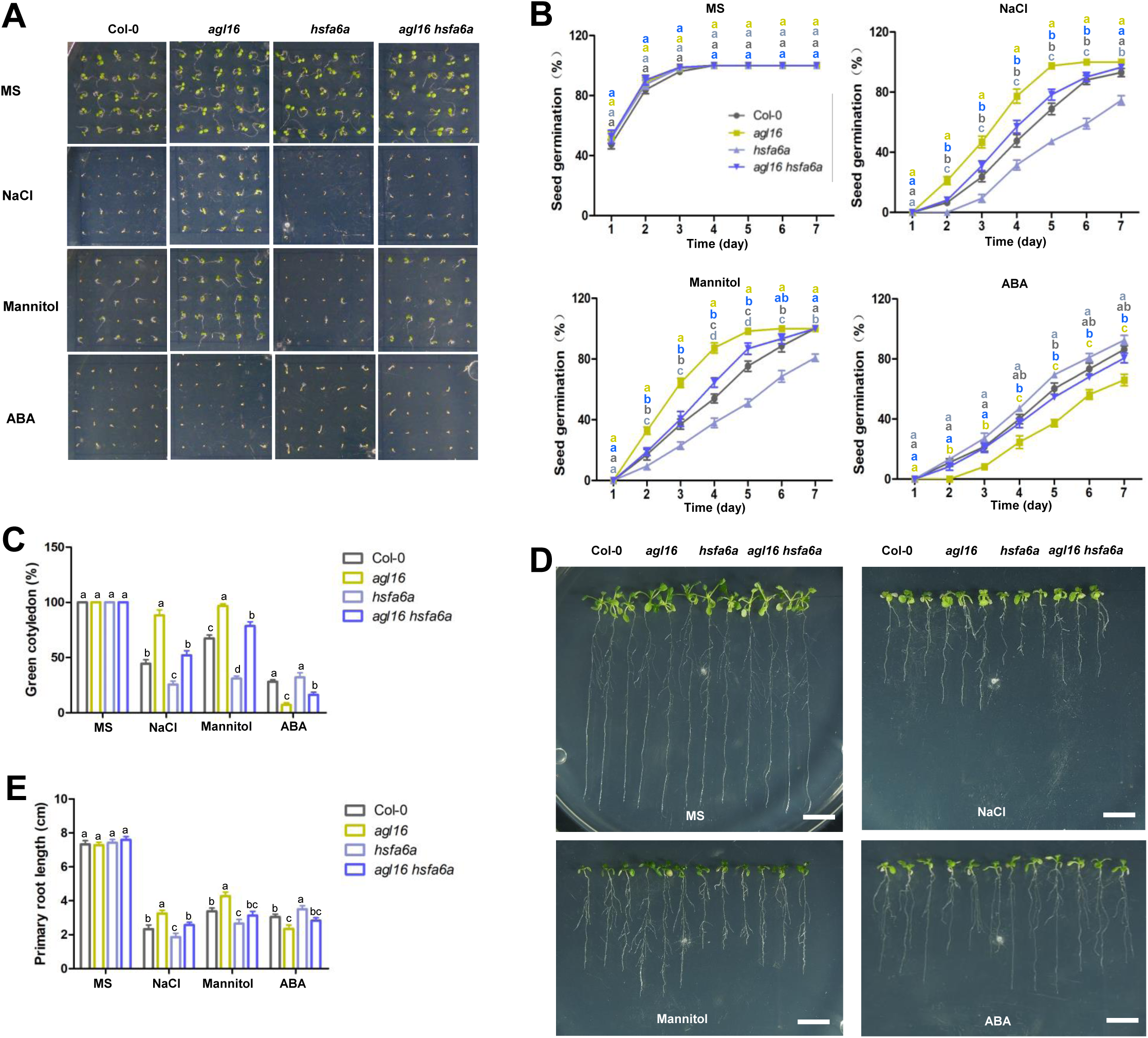
*HsfA6a* genetically acts downstream of *AGL16*. (A-C) Seed germination of Col-0, *agl16*, *hsfa6a,* and *agl16 hsfa6a* lines. Vernalized seeds were germinated on MS medium with or without 120 mM NaCl, 250 mM mannitol, or 0.5 µM ABA for 7 days before photographs were taken (A). Seed germination rate was counted at the indicated time points (B). Green cotyledon rate was counted at 7 days of germination (C). Values are mean ± SD (n=3 replicates, 60 seeds/replicate). Different letters indicate significant difference by one-way ANOVA (P < 0.05). (D-E) Primary root elongation of Col-0, *agl16*, *hsfa6a,* and *agl16 hsfa6a* lines. Seeds were germinated on MS medium for 4 days respectively, and then seedlings were transferred to MS medium with or without 120 mM NaCl, 250 mM mannitol, or 5 µM ABA and grown vertically for 7 days before photographs were taken (D) and primary root length was measured (E). Values are mean ± SD (n= 3 replicates 30 seedlings/replicate). Different letters indicate significant difference by one-way ANOVA (P < 0.05).

The *agl16 myb102* double mutant displayed an increased germination rate and green cotyledon rate compared with *myb102* mutant when subjected to mannitol and NaCl stress, while more sensitive to ABA in germination (Figure 11A-C). The primary root elongation displayed similar trends as seed germination in *agl16 myb102* mutant in response to mannitol, NaCl and ABA treatment (Figure 11D-E).

**Figure 11.**
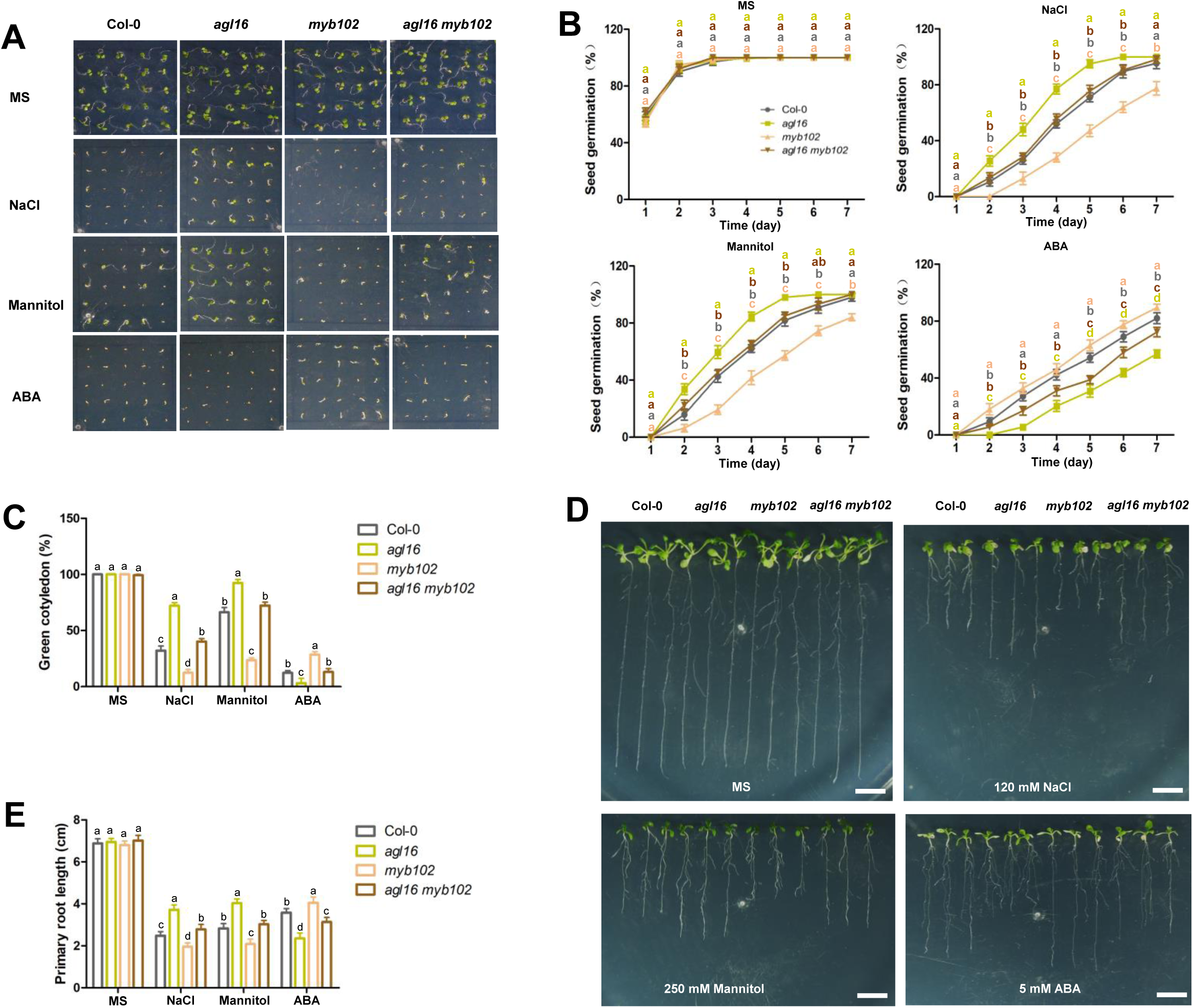
*MYB102* genetically acts downstream of *AGL16*. (A-C) Seed germination of Col-0, *agl16*, *myb102,* and *agl16 myb102* lines. Vernalized seeds were horizontally germinated on MS medium with or without 120 mM NaCl, 250 mM mannitol, or 0.5 µM ABA for 7 days before photographs were taken (A). Seed germination rate was counted at the indicated time points (B). Green cotyledon rate was counted at 7 days of germination (C). Values are mean ± SD (n=3 replicates, 60 seeds/replicate). Different letters indicate significant difference by one-way ANOVA (P < 0.05). (D-E) Primary root elongation of Col-0, *agl16*, *myb102,* and *agl16 myb102* lines. Seeds were germinated on MS medium for 4 days respectively, and then seedlings were transferred to MS medium with or without 120 mM NaCl, 250 mM mannitol, or 5 µM ABA and grown vertically for 7 days before photographs were taken (D) and primary root length was measured (E). Values are mean ± SD (n=3 replicates, 30 seedlings/replicate). Different letters indicate significant difference by one-way ANOVA (P < 0.05).

Taken together, these data support that *HsfA6a* and *MYB102* act downstream of *AGL16*, consistent with our biochemical and mocelcular evidence as the direct targets of AGL16.

## DISCUSSION

### AGL16 acts as an important negative regulator in stress response

In Arabidopsis, the MADS-box proteins play an important regulatory role in shoot development, but their roles in response to environmental stresses are still not well characterized (Yu et al., 2014a; Alvarez-Buylla et al., 2019). Previously, we reported that MADS-box TF AGL16 acts as a negative regulator in drought resistance by modulating leaf stomatal density and stomatal movement (Zhao et al., 2020). In the present study, we further demonstrated that AGL16 is also involved in ABA-mediated seed germination and seedling growth in response to salt and osmotic stress. The transcript abundance of AGL16 decreased in response to NaCl and mannitol stress while increased under ABA treatment (Figure S1B-C), indicating that AGL16 is involved in the response to multiple abiotic stresses. We found that *agl16-2* and *agl16* knockout mutants were less sensitive to NaCl and mannitol treatments during seed germination and primary root elongation but hypersensitive to ABA treatment compared with wild type. In addition, the soil-grown mutant plants exhibit enhanced the salt tolerance, while the overexpression lines show the opposite phenotypes (Figure 1-4). Together, these results clearly show that *AGL16* negatively regulates plant reponse to salt stress.

### *AGL16* is regulated by ABA signaling and responds to multiple stress and hormone signals

AGL16 belongs to the AGL17-like clade which includes ANR1, AGL21, and AGL17 members, having the overlapping expression pattern in MIKC-type group (Alvarez-Buylla et al., 2000; Burgeff et al., 2001). ANR1 and AGL21 both act as negative modulators of seed germination and by controlling the expression of *ABI3* or *ABI5* respectively (Yu et al., 2017; Lin et al., 2019). In ABA signaling pathway, ABI3, ABI4, and ABI5 act as positive regulators that are necessary for achieving growth arrest when subjected to unfavorable conditions (Giraudat et al., 1992; Finkelstein and Lynch, 2000; Yamaguchi-Shinozaki and Shinozaki, 2006). *ABI5* inhibits seed germination partly through activating the expression of LATE EMBRYOGENESIS ABUNDANT genes *EM1* and *EM6* (Finkelstein and Lynch, 2000; Carles et al., 2002; Lopez-Molina et al., 2002).

Here we found that *AGL16* expression was significantly repressed in *abi1-1*, *abi2-2*, *abi3-8* mutants under ABA treatment (Figure 5B). *ABI4*, *ABI5*, *EM6*, *RAB18*, *RD29B* were upregulated in *agl16* mutant (Figure S2), suggesting that *AGL16* is involved in ABA signaling pathway and likely acts downstream of *ABI1*, *ABI2*, and *ABI3*, but upstream or in parallel with *ABI4* and *ABI5*. Upon ABA treatment, the transcript levels of *ABA3*, *AAO3*, and *NCED3* were upregulated to modulate the level of ABA in Arabidopsis (Chan, 2012; Vishal and Kumar, 2018). In *agl16* mutant, the expression level of *ABA2*, *NCED3*, *AAO3* was elevated (Figure S2M, O-P), likely contributing to the hypersensitivity to exogenous ABA in *agl16* mutant. Several TFs, such as ABR1, RGL2, WRKY2, WRKY41 were reported to regulate seed germination depending not only on ABA signaling pathway, but also on ABA biosynthesis (Pandey et al., 2005; Piskurewicz et al., 2008; Jiang and Yu, 2009; Ding et al., 2014).

Besides, a few studies reported that salt stress induces expression of genes related to auxin biosynthesis and transport, such as *nitrilase 1* (*NIT1*) and *PIN2* (Bao and Li, 2002; Sun et al., 2008). Ethylene and auxin have been shown to regulate growth and development processes, including primary root elongation (Swarup et al., 2007). In this study, *AGL16* responds to multiple stresses and hormones ethylene and IAA (Figure S1D) and affects cell division (Figure 3), suggesting that AGL16 may serve as a node integrating phytohormone signals derived from environmental stresses to mediate the adaptive response of primary root development.

### AGL16 enhances salinity tolerance by modulating ion homeostasis

High salinity can cause hyperosmotic stress and sodium (Na^+^) toxicity in plants. Maintaining Na^+^ and K^+^ homeostasis in the cytosol by inhibition of Na^+^ influx and promotion of K^+^ uptake is critical for plant survival in salt tolerance (Yokoi et al., 2002; Sunarpi et al., 2005; Ma et al., 2011). Lower levels of Na^+^ were accumulated in the shoots of *agl16* mutant compared with Col-0 when exposed to NaCl (Figure 4D), while no significant difference was observed in roots (Figure 4F), indicating Na^+^ level was decreased in shoots by increasing Na^+^ influx into roots mostly via enhanced the expression of *HKT1;1*. *AtHKT1;1* is expressed mainly in the root xylem parenchyma cells and plays an essential role in Na^+^ exclusion from leaves and K^+^ homeostasis in leaves under salinity stress (Maser et al., 2002; Horie et al., 2008). *AGL16* was highly expressed in the stele (Figure S1Ah), which overlaps with *HKT1;1* expression in the root, supporting that AGL16 regulates *HKT1;1* expression. It has been reported that ABI4, represses *HKT1;1* expression in the root stele and alters Na^+^ unloading from xylem vessels to xylem parenchyma cells, increasing salt tolerance in *abi4* mutants (Shkolnik-Inbar et al., 2013). Salt stress-induced bZIP24 regulates plant salt tolerance by directly or indirectly repressing the expression of *HKT1;1* (Yang et al., 2009). *AtNHX1* was not significantly affected by AGL16 in response to salt stress (Figure S3). Therefore, The loss-of-*AGL16* improves salinity tolerance by enhancing the capacity of Na^+^ extrusion rather than Na^+^ sequestration.

### AGL16 negatively regulates a large array of stress-responsive genes

Comparative transcriptomic analyses revealed that many DEGs involved in the response to heat, wounding, and jasmonic acid were found preferentially enriched in KO and OX lines under normal condition (Figure S5A-B), indicating that AGL16 may play an important role in the response to abiotic stresses and hormone signals. A number of genes related to water deprivation, salt stress, abscisic acid, osmotic stress were significantly enriched in KO vs WT (salt stressed) group (Figure S5C), indicating that the enhanced salt tolerance of *agl16* may be attributed to increased ion and osmotic signaling, which activates the expression of downsteam stress-responsive genes, for instance, the stress- or ABA-responsive genes showed in Figure 5C, which are known to participate in abiotic stress response and improve plant resistance to multiple stresses when overexpressed (Nakashima et al., 2000; Zhu, 2002; Bartels and Sunkar, 2005).

TFs are major regulators of gene expression and also involved in the crosstalk between plant hormones and stress signaling (Fujita et al., 2006; Ku et al., 2018; Xie et al., 2019). Several stress-responsive TF genes were enriched in our RNA-seq data (Figure 6C), including *HsfA6a* and *MYB102*. *HsfA6a* (Hwang et al., 2014) and *MYB102* (Denekamp and Smeekens, 2003) are known to be induced by abiotic stress or ABA. *AtHsfA6a* is highly induced by exogenous ABA, NaCl, and drought. The overexpression lines of *HsfA6a* are hypersensitive to ABA and exhibit enhanced tolerance against salt and drought stress (Hwang et al., 2014).

MYB proteins are notable for their roles in regulation of multiple responses to hormone signaling, biotic and abiotic stresses (Dubos et al., 2010; Roy, 2016). *MYB102* is expressed in almost all organs and rapidly induced by osmotic stress and ABA (Denekamp and Smeekens, 2003; Zhu et al., 2018), overlapping with the expression profiles of *AGL16*, which suggests that it may play an important role in abiotic stress resistance. Indeed, loss-of-*MYB102* significantly impairs the resistance to salt and osmotic stress in germination and root elongation (Figure 11).

Our genetic and biochemical analyses demonstrate that *AGL16* acts upstream of *HsfA6a* and *MYB102* and AGL16 suppresses the transcription of *HsfA6a* and *MYB102* by directly binding to their promoter (Figure 8-11).

### AGL16 acts as a negative regulator of stress responses to balance between growth and stress response

Sensile plants live in a constantly fluctuating environment. They must equipped with highly efficient and effective mechanisms to balance between growth and response to stresses. Plants maximize growth under favorable conditions, while under environmental stresses, they allocate resources to cope with stresses and slow down or stop growth. This dynamic switch must be finely tuned to ensure plant success in nature. In this study, we have characterized the MADS-box TF AGL16 as an important negative regulator that keeps stress response off or down when environmental stresses are absent or low. AGL16 regulates plant stress tolerance by transcriptionally suppressing a large array of stress responsive target genes including *HKT1;1*, *HsfA6a,* and *MYB102* to maintain ion homeostasis and proper response to stress. Furthermore, AGL16 may mediates ABA signaling by negatively regulating ABI5 and ABA content. Therefore, our study suggests that AGL16 serves as an important modulator balancing between plant response to stress and plant growth, and provide a beneficial candidate for improving crop stress resistance.

## METHODS

### Plant materials and growth conditions

The wild type used in our study was *Arabidopsis thaliana* ecotype Columbia-0 (Col-0). All mutants and transgenic plants used in our work are in the Col-0 background. A homozygous *AGL16* loss-function mutant *agl16* (Salk_104701) was ordered from Arabidopsis Biological Resource Center (ABRC). The CRISPR/Cas9-edited *AGL16* knockout mutant *agl16-2*, *35S:AGL16*, *35S:HA-AGL16*, AGL16*pro: GUS*, *35S:AGL16-GFP* transgenic plants were obtained by transforming into Col-0 via *Agrobacterium* (C58C1) transformation using the Arabidopsis floral-dip method (Clough and Bent, 1998). *agl16 CyclinB1:1*, *agl16-2 CyclinB1:1*, *35S:AGL16 CyclinB1:1*, *agl16 hsfa6a* and *agl16 myb102* lines were obtained by genetic crossing separately.

Arabidopsis seeds were surface sterilized with 10% bleach for 15-20 minutes, and then washed four times at least with distilled water. Seeds were vernalized at 4°C for 2 days in darkness, and then horizontally or vertically germinated on MS (Murashige and Skoog) medium with 1% (w/v) sucrose at 22 °C, 60-80% relative humidity under 16-h light/8-h dark cycles. For seed germination assay, seeds were horizontally germinated on MS mediums with or without 120 mM NaCl, 250 mM Mannitol, and 0.5 µM ABA for 7 days, germination (emergence of radicles) and post-germination growth (green cotyledon appearance) were scored at the indicated time points. For analysis of PR length, 4-day-old seedlings were transferred to MS mediums with or without 120 mM NaCl, 250 mM Mannitol, and 5 µM ABA for 8 days and grown vertically, the PR length of seedlings and fresh weights were measured at the indicated time points.

### Analysis of GUS activity

The transgenic lines *AGL16pro:GUS* were obtained by the 2.5 kb promoter of *AGL16* was cloned into pCB308R vector (Lei et al., 2007). The different tissues of seedlings and adult seedlings of *AGL16pro: GUS* transgenic plants were stained as described previously (Jefferson et al., 1987). GUS staining solution was prepared as described before (Cai et al., 2014). The GUS activities of individual parts were observed using a light microscope with a camera (HiROX).

### Subcellular localization

The *AGL16*-coding region was constructed into pGWB5 vector (Nakagawa et al., 2007) and obtained transgenic plants by *Agrobacterium* (C58C1) transformation using the Arabidopsis floral-dip method. The green fluorescence of *35S:AGL16-GFP* transgenic root tissues were observed under a ZEISS880 confocal laser scanning microscope with an excitation of 488 nm and an emission of 525 nm condition.

### RT-PCR and qRT-PCR analysis

Total RNA of different tissues as indicated using TRIzol reagent (Invitrogen, Carlsbad, USA). RNA reverse reaction was carried out using Prime Script RT reagent kit (Takara, Dalian, China). The reverse reaction products were used as templates for RT-PCR amplification and qRT-PCR analysis. Applied Biosystem StepOne real-time PCR system and SYBR Premix Ex Taq II kit (Takara, Dalian, China) were used for the qRT-PCR detection. The transcript levels of different genes were normalized to that of UBIQUITIN5 (UBQ5) in qRT-PCR analysis.

### RNA-seq analysis

For RNA-seq analysis, the *Arabidopsis* roots from 7-day-old Col-0, *agl16*, and OX-17 seedlings exposed to medium with salt treatment for 0, 3 hours were respectively collected, then RNA was isolated using RNAprep Pure Plant Kit (TIANGEN, Beijing, China). The total RNA integrity was evaluated by Bioanalyzer 2100 (Agilent, California, USA) (RIN ≥ 7, 28S/18S ≥ 1.5), and RNA-seq library was sequenced by Beijing Biomarker Technology Company. In RNA-seq data, differentially expressed genes (DEGs) were characterized by the threshold (the absolute value of Log_2_ (fold change) ≥ 1.5, FDR ≤ 0.05). GO term enrichment analysis for each differentially expressed genes was conducted by David tools (https://david.ncifcrf.gov/).

### ChIP assay

7-day-old *35S:HA-AGL16* transgenic seedlings and Col-0 plants were used for ChIP assay. The procedure was conducted as described previously (Cai et al., 2014). The purified DNA fragments were used as templates for PCR amplification and qRT-PCR detection with the same primers. For PCR analysis, PCR products were detected by 2% agarose gel electrophoresis. The DNA binding ratio was calculated by qRT-PCR (Mukhopadhyay et al., 2008). β-tubulin8 was used as a negative control.

### Transient expression assay

Transient expression assay was carried out as described previously (Wang et al., 2016). The *AGL16*-coding region was constructed into pGreenII 62-SK to generate an effector, and about 2000 bp promoter of *HKT1;1*, *HsfA6a*, *MYB102* or putative 26 bp fragment containing CArG motifs from these target genes promoter were respectively fused to pGreenII 0800-LUC to generate reporters (Hellens et al., 2005). The effector and reporter were subsequently transfected into *Arabidopsis* mesophyll protoplasts from 4-week-old wild-type (Col-0) leaves in short light as described (Yoo et al., 2007). The firefly LUC and REN activities luciferase were detected by transient dual-luciferase reporter assay system (Promega, Madison, USA), and LUC/REN ratio was calculated. The REN activity was used as an internal control.

### Yeast one-hybrid assay

Yeast one-hybrid assay was conducted as described previously (Mao et al., 2016). The *AGL16*-coding region was constructed into pAD-GAL4-2.1 (pAD) and the putative 26 bp fragment containing CArG motifs from *HKT1;1*, *HsfA6a*, *MYB102* promoter were respectively cloned into pHIS2 (pBD) (Mu et al., 2009). These constructs of pAD and pHIS2 plasmids were introduced into Y187 yeast strain and grown on SD/-Trp-Leu medium with or without His plus 20 mM 3-aminotriazole and diluted as indicated (1:1, 1:10, 1:100, 1:1000).

### Electrophoretic mobility shift assay

Electrophoretic mobility shift assay (EMSA) was performed as described previously (Miao et al., 2018). The *AGL16*-coding region was cloned into pET28a vector and His-AGL16 fusion protein was expressed in E. coli Rosseta2 strain. 30 bp free probes containing CArG motifs, non-competitor probes (mutated) were synthesized with biotinlabeled at the 5′ end by a commercial company (Sangon Biotech, Shanghai, China), while competitor probes did not. Complementary ssDNA probes were mixed, 95 °C for 10 min and slowly cooled down to 25 °C. EMSA was performed using a LightShift™ EMSA Optimization and Control Kit (Thermo Fisher Scientific, Waltham, USA). For competition assays, different concentrations of competitor and non-competitor probes were added into the reaction system. Each reaction was loaded on a 6% native polyacrylamide gel in 0.5 × TBE buffer to electrophoresis. The results were detected using a CCD camera system (Image Quant LAS 4000).

### Genetic analysis

The *agl16* loss-of-function mutant was a female parent, *abi5*, *hsfa6a* or *myb102* loss-of-function mutant was as male parent. They were crossed by transferring mature anthers of male parent to stigmas of female parent and obtained the homozygotes of double mutants.

### Statistical analysis

Statistical analysis was conducted using one-way ANOVA. Values are the mean ± SD and P<0.05 was considered statistically significant. Different letters indicate a significant difference.

### Accession numbers

Sequence data from this study can be found in the GenBank/EMBL libraries under the following accession numbers: *AGL16* (AT3g57230), *UBQ5* (AT3g62250), *ABA1* (AT5g67030), *ABA2* (AT1g52340), *ABA3* (AT1g16540), *NCED3* (AT3g14440), *AAO3* (AT2g27150), *CYP707A1* (AT4g19230), *CYP707A2* (AT2g29090), *CYP707A3* (AT5g45340), *CYP707A4* (AT3g19270), *ABI1* (AT4g26080), *ABI2* (AT5g57050), *SnRK2.2* (At3g50500), *SnRK2.3* (At5g66880), *ABI3* (AT3g24650), *ABI4* (AT2g40220), *ABI5* (AT2g36270), *EM1* (AT3g51810), *EM6* (AT2g40170), *RAB18* (AT5g66400), *RD29B* (At5g52300), *RD22* (At5g25610), *RD29A* (AT5g52310), *CBF4* (AT5g51990), *SOS1* (AT2g01980), *SOS2* (AT5g35410), *NHX1* (AT5g27150), *HKT1;1* (AT4g10310), *HsfA6a* (At5g43840), *MYB102* (At4g21440), *COR15B* (AT2g42530), *NAC055* (AT3g15500), *bZIP1* (AT5g49450), *RAV1* (AT1g13260), *NHX2* (AT3g05030), *HAK5* (AT4g13420), *RAP2.6* (AT1g43160), *MYC5* (AT5g46830), *RAS1* (AT1g09950).

## Supplemental information

Figure S1. Expression pattern of AGL16 and response to abiotic stresses during root development.

Figure S2. The expression levels of ABA signaling, biosynthesis and catabolic-related genes in Col-0, *agl16*, and OX-17 lines.

Figure S3. Expression profiles of salt stress-responsive genes in Col-0, *agl16*, OX lines.

Figure S4. The distribution of differentially expressed genes (DEGs) in KO vs WT-control, OX vs WT-control, KO vs WT-salt, and OX vs WT-salt groups.

Figure S5. GO term enrichment analysis of genes affected by AGL16.

Figure S6. The transcript levels of stress-responsive genes of RNA-Seq microarray analysis.

Table S1. List of overlapped genes related to salt stress between KO vs WT-control, OX vs WT-control, KO vs WT-salt, and OX vs WT-salt groups.

Table S2. Primers used in this study.

## ACKNOWLEDGEMENTS

This study was supported by grants from National Natural Science Foundation of China (31900230 to P.Z.), China Postdoctoral Science Foundation (2020T130634 and 2019M652200 to P.Z.) and Youth Innovation Foundation of University of Science and Technology of China (WK2070000186). We thank the ABRC for providing the mutant seeds.

## AUTHOR CONTRIBUTIONS

P.Z. and C.X. designed the experiments. P.Z., J.Z., Y.C., J.W., J.X., L.S., S.M. performed the experiments and data analyses. P.Z. wrote the manuscript. C.X supervised the project and revised the manuscript.

